# An experimentally-derived measure of inter-replicate variation in reference samples: the same-same permutation methodology

**DOI:** 10.1101/797217

**Authors:** David C. Handler, Paul A. Haynes

## Abstract

The multiple testing problem is a well-known statistical stumbling block in high-throughput data analysis, where large scale repetition of statistical methods introduces unwanted noise into the results. While approaches exist to overcome the multiple testing problem, these methods focus on theoretical statistical clarification rather than incorporating experimentally-derived measures to ensure appropriately tailored analysis parameters. Here, we introduce a method for estimating inter-replicate variability in reference samples for a quantitative proteomics experiment using permutation analysis. This can function as a modulator to multiple testing corrections such as the Benjamini-Hochberg ordered Q value test. We refer to this as a ‘same-same’ analysis, since this method incorporates the use of six biological replicates of the reference sample and determines, through non-redundant triplet pairwise comparisons, the level of quantitative noise inherent within the system. The method can be used to produce an experiment-specific Q value cut-off that achieves a specified false discovery rate at the quantitation level, such as 1%. The same-same method is applicable to any experimental set that incorporates six replicates of a reference sample. To facilitate access to this approach, we have developed a same-same analysis R module that is freely available and ready to use via the internet.

## 1. Introduction

Shotgun proteomics experiments that seek to compare ‘reference’ and ‘treated’ states of a given sample will often contain thousands of individual comparisons, each requiring statistical validation. In such circumstances, repeated use of the Student’s t-test will invariably introduce false discoveries into the results. A Student’s t-test with a significance cut-off threshold P value of 0.05 produces 95% confidence i.e. 5% of tests will have an equal likelihood to be attributable to chance instead of experimental factors [1-4]. This is a follow-on conclusion from the probability that at least one test in an experiment will be significant, which is described as 1 – (1 - α)^k^, where α is the significance cut-off and k is the number of tests conducted [5].

Multiple Testing Corrections (MTCs) were introduced to help address this limitation [6], including the use of Q values rather than P values [7], Bonferroni correction [8,9], Benjamini-Hochberg (BH) adjusted t-test [10], Bonferroni-Holm test [11], and the Benjamini-Yoav (BY) test [12]. MTCs, however, are often overly conservative and increase the false negative rate by eliminating otherwise valid protein identifications. This is especially a problem at the protein-quantitation level; MTCs, by their nature, contribute to a lessening of the protein quantitation false discovery rate (PQ-FDR) at the expense of otherwise valid protein identifications [13,14]. In the context of this article, PQ-FDR is defined as false discoveries arising from comparative quantitative proteomics calculations between one or more samples. There is always a balance to be struck between stringency and accuracy when controlling false discoveries at the protein quantitation level [15]. A recent study in this area applied Bayesian statistics to great effect, detecting a greater number of relevant protein quantitation changes in previously published data sets [16]. There are also numerous software packages available which incorporate various other MTC approaches, including Proteus [17], DAPAR and ProStaR [18], MSqRob [19], UbiA-MS [20], ProteoSign [21], msVolcano [22], FDRtool [23], MSstats [24], and limma [25].

Although the use of MTC correction methods in the proteomics field is not standardized [9,26-28], MTCs are an important tool that researchers can employ for extracting the best results from their dataset i.e. finding the balance between reducing noise without losing signal. This desire to reduce the noise in the system led us to ask the question: is there a better way to quantify variability between replicate analyses of a reference sample ?

One established approach for assessing variability across a sample set is to use permutation analysis, based upon the Significance Analysis of Micro-array (SAM) permutation methodology [29,30]. This is a similar theoretical framework to that used for permutation analysis within Perseus [31], a well-established data analysis program in the MaxQuant environment [32]. The SAM permutation analysis method assigns a score to each gene on the basis of change in gene expression relative to the standard deviation of repeated measurements. SAM then uses redundant permutations of repeated measurements to estimate the percentage of genes identified by random chance as an artefact of the method, which is used to calculate the false discovery rate. The permutations are performed across all of the ‘reference’ and ‘treated’ sample replicates within a given experimental data set. Those genes with scores higher than the specified threshold are deemed potentially significant, and the threshold can be adjusted to identify smaller or larger sets of genes, with FDR calculated for each set.

The same – same method introduced in this study, in contrast, employs non-redundant permutations of experimentally repeated measurements of protein abundance in replicate analysis of a defined reference sample. The permutations are performed on data from the reference samples only, isolated from the ‘treated’ samples. This is used to generate a single average Q value indicative of the degree of variation of abundance across the reference sample replicates. Proteins which reach a defined statistical significance threshold are deemed to be false discoveries at the protein quantitation level, since comparing a reference sample against itself should theoretically yield no changes in protein abundance. It is important to emphasize that the underlying assumption is that the biological variability between reference samples is zero, so this approach is accounting for the technical variability. This facilitates subsequent assessment of induced biological variation between reference samples and treated samples.

A specified false discovery rate in the same – same analysis of replicates of the reference sample is used to generate a Q value threshold, and that value can then be carried forward to the subsequent analysis of a reference sample versus ‘treated’ sample within the same larger experimental data set. One of main the applications of this method for determining an experimentally-derived measure of reference sample variability is that it can subsequently be used to modify an existing MTC protocol for downstream analysis, thus minimising the PQ-FDR without introducing false negatives. By performing a specific permutation analysis to measure the variability inherent within reference sample replicates, we can produce an experimentally modulated Q value threshold for use with MTCs when comparing the reference sample to treated samples. In essence, rather than using a default Q value of .05, or choosing a more stringent value, we are employing a Q value threshold that is experimentally determined for each set of samples analyzed. The same-same method represents another tool in the proteomics toolbox, and can be used to enable the extraction of additional biological knowledge from large-scale datasets.

## 2. Materials and Methods

### 2.1 Label free quantitative proteomics data sets

To demonstrate the utility of the same-same approach we reanalyzed two sets of previously published label free quantitative shotgun proteomics data. Protein identification and Normalized Spectral Abundance Factor (NSAF) values [33,34] were sourced from previously published studies from our laboratory on cultured Cabernet Sauvignon grape cells grown at different temperatures [35] (ProteomeXchange identifier PXD000977) and leaf tissue of IAC1131 rice plants exposed to drought stress [36] (ProteomeXchange identifier PXD004096). The cultured Cabernet Sauvignon grape cell data consists of six biological replicates of cell cultures maintained at 26°C, as the optimum, or control, temperature, and biological triplicate analysis of cells maintained at 18°C and 10°C as moderate and extreme cold stress conditions, and 34°C and 42°C as moderate and extreme heat stress conditions. The rice leaf data consists of two sets of three biological replicates each of unstressed plants as controls, and biological triplicate analysis of plants exposed to moderate drought stress, extreme drought stress, and extreme drought stress followed by recovery.

### 2.2 Same – same permutation analysis of reference samples

For analysis using the same-same workflow, six replicates of a reference sample are run through a PSM (peptide-to-spectrum matching) engine such as GPM or ProteomeDiscoverer and protein identification lists are exported as csv files. At a minimum, protein identifier and peptide count are needed. Next, these six replicates are grouped into two sets of triplicates by the use of inner joins (dummy state ‘control’, dummy state ‘treatment’), and a test array is formed through a full join of the states [37]. One hundred Student’s t-tests are conducted on spectral counts from each identified protein comparison with significance cut-off values from 0.01 to 1, stepped at 0.01 intervals. All proteins found at different quantitation levels are considered false discoveries, since comparison between two data sets of the same sample type would theoretically give identical quantitation with no observed changes. This process is repeated for all ten combinations of non-redundant triplet pairs that six replicates can form. The MTC analysis then begins by iterating over this 10x t-test array and applying one of five user-specified MTC methods (BH, Benjamini-Yoav, Bonferroni, Hommel [38], and Bioconductor Q [39]. The program then averages the MTC test results from all arrays examined, and reports the point at which the significance cut-off corresponds to a user-specified PQ-FDR.

The same-same methodology is automated through an R script. Source code is available from https://bitbucket.org/peptidewitch/samesame/, and a freely accessible working web version can be found at https://peptidewitch.shinyapps.io/samesame. The R Shiny web-app provides three distinct outputs from the same-same analysis:

1. A series of Q value vs FDR bar plots (x axis 0.01 to 1, stepped at 0.01) from all ten triplet paired combinations,
2. A series of P value histograms of these same combinations, and,
3. A numerical value that corresponds to the user-specified MTC cut-off that produces the desired PQ-FDR (default 1%).

Input data types are not constrained to spectral counts, as in theory any data type that consists of protein identifications coupled with abundance or intensity value measurements can be used. However, the first generation of the analysis tool was designed and tested using spectral counting-based data, so it is recommended that spectral counts or spectral abundance factors be used initially.

### 2.3 Perseus permutation analysis of reference samples

To serve as a comparison against the same-same process, the same NSAF data from both Grape and Rice samples as were reanalyzed using Perseus software [31]. Spreadsheet files containing NSAF values for each set of samples were uploaded to Perseus through a generic matrix upload. Using the two-sample module, we applied the Perseus permutation method as a form of truncation using ungrouped (no grouping preserved), 250-count permutation analysis on two-tailed Student’s t-testing arrays with BH correction, comparing six reference replicates with three replicates from each of the ‘treated’ sample states, with the specified FDR thresholds ranging from 1-5%.

## 3. Results

The following section details how same-same approach was applied to the published data from Grape and Rice cells. Figure 1 displays the outputs described above for the same-same analysis conducted on the grape cell culture label-free data when specifying BH correction and 1% PQ-FDR. The end-point of the same-same process is the modulated Q value, in this example 0.054 (Figure 1C), produced from averaging the threshold values in Figure 1A at the desired PQ-FDR value. This value can be used for downstream analysis on subsequent control vs treatment samples as a modulator for the chosen MTC.

**Figure 1.**
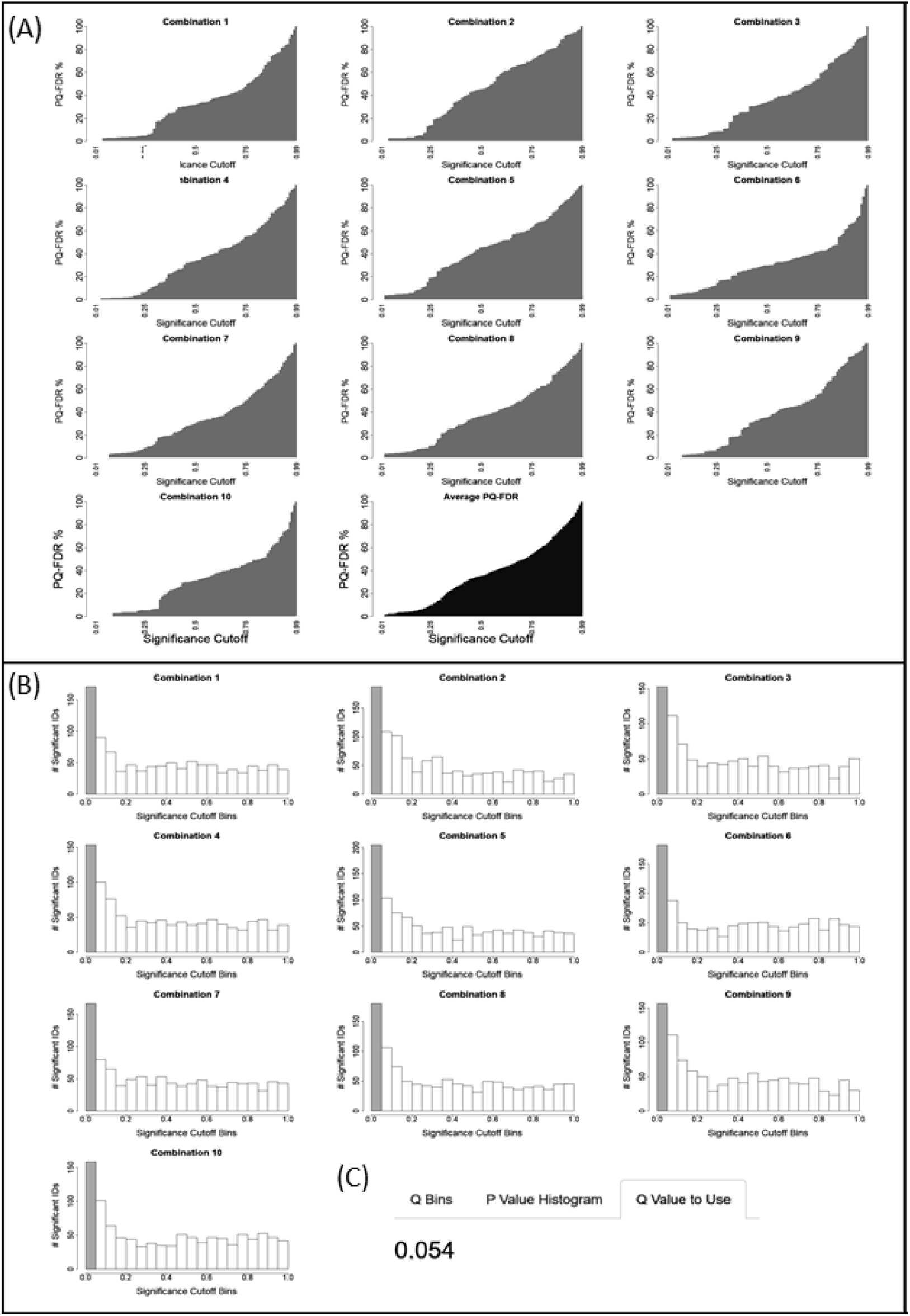
Screenshots from the same/same shiny apps module (https://peptidewitch.shinyapps.io/SameSame), using the grape cell control samples and specifying BH correction at 1% PQ-FDR. (A) Q value vs PQ-FDR bar plots (x axis 0.01 to 1, stepped at 0.01) for all ten triplet paired permutations generated from six replicate analyses of a reference sample (see Figure 1) (B) P value histograms for each permutation, showing number of significantly expressed protein identifications sorted into P values bins in increments of 0.05. (C) displays a single numerical value which produces the desired PQ-FDR value (default BH at 1%, can be user specified).

Figure 2 presents the subsequent downstream analysis of the grape cell cultures grown at different temperatures. Figure 2A shows the number of proteins found to be significantly differentially expressed in terms of protein fold change when comparing the set of six reference replicates to the set of three replicates of cells grown at each temperature. These are analysed using different statistical measures of significance: P values of 0.05 and 0.01 for a student’s t-test, BH Q value of 0.05, and BH using the same-same derived Q value (SS-Q), and specifying PQ – FDR of 1%, 2% or 3%. It is evident that the same-same derived Q values at 1% PQ-FDR produce results very similar to the use of default BH Q values, which is expected given that the SS-Q value used is very close to the 0.05 BH-Q value threshold. The two approaches give similar results, although it is noticeable that at a specified PQ – FDR of 3%, the comparison with the largest effect size (Figure 2E) shows significantly more differentially expressed proteins.

**Figure 2.**
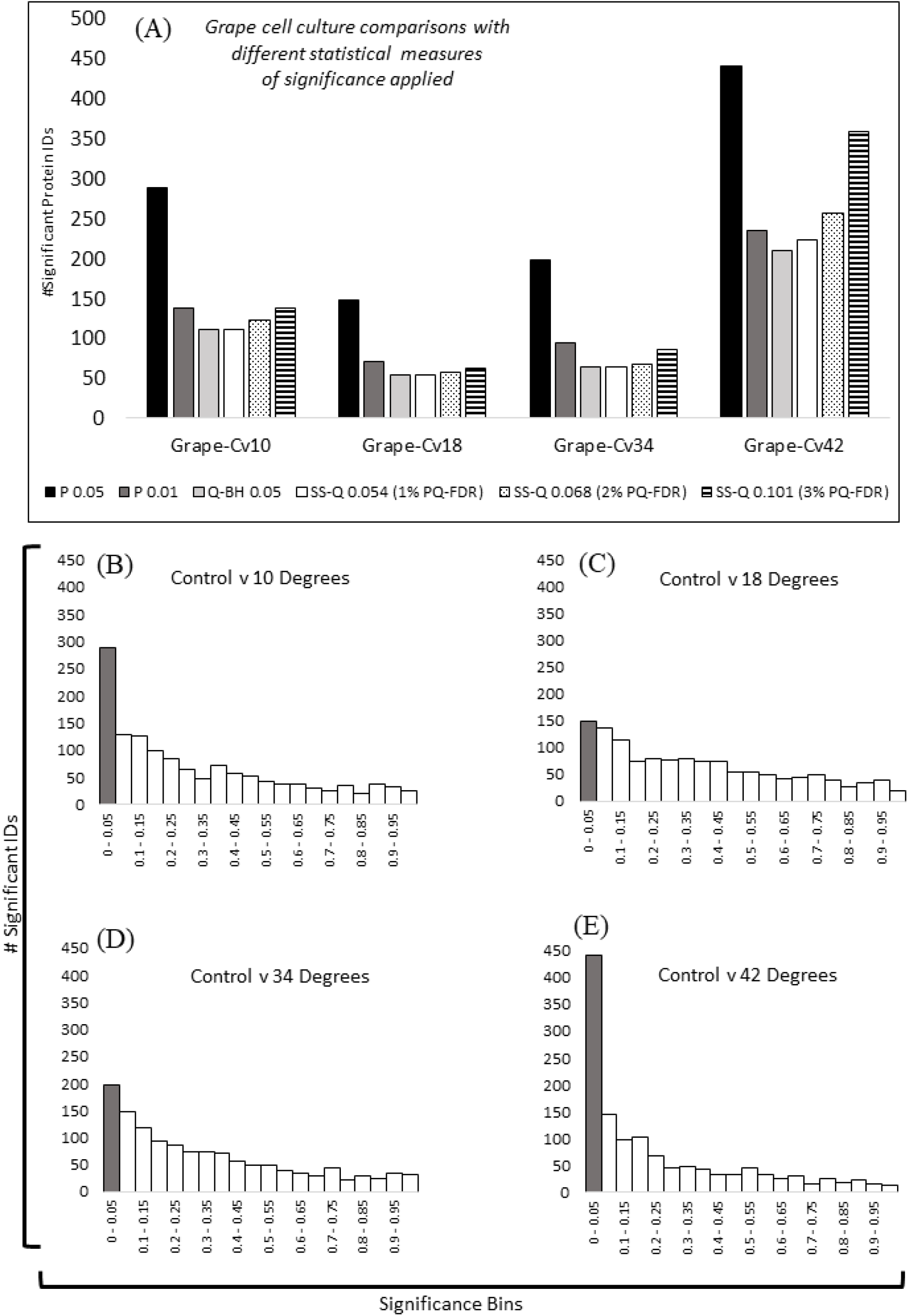
Grape cell culture comparisons with application of different statistical significance measures. Cells grown at 26°C were designated as the reference sample, and compared with cells grown at 18°C (moderate cold), 10°C (extreme cold), 34°C (moderate heat), and 42°C (extreme heat). Panel A displays the number of significantly differentially expressed protein identifications found for each comparison using P values at .05 and .01, Benjamini-Hochberg adjusted values at 0.05, and BH using the same-same derived Q value (SS-Q), and specifying PQ – FDR of 1%, 2% or 3%. Panels B to E contain P value histograms showing the number of significantly expressed protein identifications sorted into P value bins in increments of 0.05, for each of the four experimental comparisons performed, as indicated.

Figure 3 presents the same type of analyses as shown in Figure 1 for the data derived from comparative analysis of leaf tissue from IAC1131 rice plants exposed to different levels of drought stress. Interestingly, in contrast to figure 2, it is clear that in this case there is a direct correlation between observed effect size and number of differentially expressed proteins identified using the SS-Q approach. In comparisons with greater effect size as observed in P value histograms (Figure 3E,3F,3G), the same-same derived Q Values are able to identify a greater number of differentially expressed proteins than were found using the default BH Q value, and at 3% PQ-FDR are approaching the number of differentially expressed proteins found using uncorrected P values.

**Figure 3.**
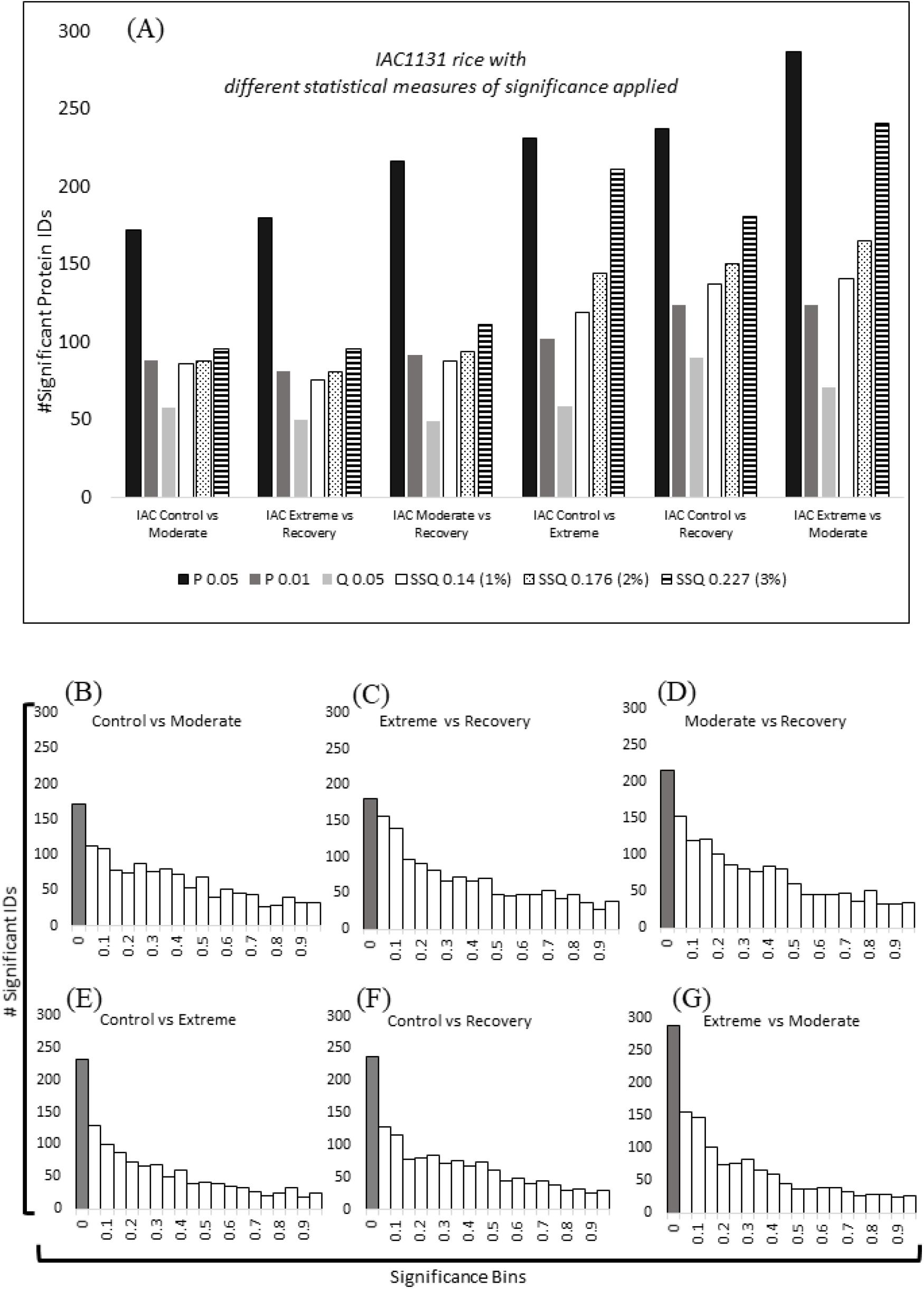
IAC1131 rice samples drought stress comparisons with application of different statistical significance measures. Control plants were unstressed, and compared with plants exposed to moderate drought stress, extreme drought stress, or extreme drought stress followed by recovery. Panel A displays the number of significantly differentially expressed protein identifications found for each comparison using P values at .05 and .01, Benjamini-Hochberg adjusted values at 0.05, and BH using the same-same derived Q value (SS-Q), and specifying PQ – FDR of 1%, 2% or 3%. Panels B to G are P value histograms showing the number of significantly expressed protein identifications sorted into P value bins in increments of 0.05, for each of the six experimental comparisons performed, as indicated.

Table 1 presents the results of analyzing the grape and rice cell NSAF data referred to above using different analysis approaches, including Student t-tests with and without BH correction, application of same – same derived Q values to a BH corrected t-test at specified PQ-FDR values ranging from 1% to 5%, and t-tests using Perseus permutations at specified PQ-FDR values ranging from 1% to 5%. The table shows the number of proteins which are reported to be significantly differentially expressed when comparing the reference samples against the grape cells grown at four different temperatures, and the rice cells grown under three different watering regimes. It is clear from these comparisons that, as expected, the uncorrected student’s t-test gives a much greater number than any sort of correction. The BH correction reduces the number of significant proteins by approximately 95%. The Perseus permutation processing is even more strict and, for example, produces zero significant identifiers in more than half of the grape sample comparisons. In contrast, the same-same-modulated BH test is able to detect significantly differentially regulated proteins for every test case for both tissue types while always remaining well below the results reported from uncorrected Student’s t-testing P values. Multiple testing correction still takes place, but the experimentally derived Q value thresholds allow for the recovery of a greater number of significant differences at the protein quantitation level.

**Table 1.** **Comparison of number of protein identifications retained using different analysis approaches to assess protein quantitation false discovery rate**

| Comparisons | t-test<sup>a</sup> |  | Perseus permutations BH |  |  |  |  | Same-Same BH |  |  |  |  |
| --- | --- | --- | --- | --- | --- | --- | --- | --- | --- | --- | --- | --- |
| **Grape** | **.05** | **BH** | **1%<sup>b</sup>** | **2%** | **3%** | **4%** | **5%** | **1%** | **2%** | **3%** | **4%** | **5%** |
| Cont vs 10 C | 735 | 42 | 0 | 0 | 0 | 0 | 0 | 48 | 54 | 72 | 91 | 99 |
| Cont vs 18 C | 560 | 29 | 0 | 0 | 0 | 9 | 9 | 29 | 34 | 43 | 51 | 65 |
| Cont vs 34 C | 432 | 21 | 0 | 0 | 0 | 0 | 6 | 24 | 28 | 36 | 42 | 48 |
| Cont vs 42 C | 775 | 83 | 0 | 0 | 0 | 0 | 44 | 108 | 116 | 133 | 166 | 199 |
| **Rice** | **.05** | **BH** | **1%** | **2%** | **3%** | **4%** | **5%** | **1%** | **2%** | **3%** | **4%** | **5%** |
| Cont vs Ext | 543 | 12 | 0 | 0 | 0 | 0 | 0 | 15 | 16 | 17 | 17 | 17 |
| Cont vs Mod | 481 | 13 | 0 | 0 | 0 | 0 | 0 | 13 | 17 | 20 | 20 | 20 |
| Cont vs Recov | 568 | 21 | 0 | 0 | 0 | 0 | 0 | 22 | 33 | 40 | 40 | 40 |

<sup>a</sup> .05 = standard 2-sample t-test, BH = Benjamin-Hochberg corrected 2 sample t-test
<sup>b</sup> protein quantitation false discovery rate assessed at 1%-5% using the approaches indicated

## 4. Discussion

The correlation observed between effect size and number of differentially expressed proteins found in the dataset presented in Figure 3 has also been found in numerous other datasets we have analysed. In general, SS-Q values are generally better suited to those datasets that show a larger effect size. This may be due to the fact that not all quantitatively different proteins in a small effect sample are false positives, or may be a consequence of NSAFs overstating expression change ratios for protein identifications based on lower spectral counts, which can help to increase the effect size [4]. While the use of higher Q value thresholds raises the implicit question of whether or not the dataset contains too much noise, it is important to remember why the same-same experiment is conducted in the first place. If, in an experiment where we expect there to be minimal noise, we demonstrate that there is a SS-Q threshold value that produces 1% PQ-FDR between sets of control or reference replicates, then in a closely related experiment with the same reference sample using the same threshold value, we can infer experimentally that the specified PQ-FDR has been achieved.

It is important to stress, however, that this method is suited more towards initial discovery, and that follow-up experimentation must employ orthogonal validation protocols. In order to obtain an experimentally-derived PQ-FDR of 1%, or other specified value, the same-same method is a very useful tool, because inferring the PQ-FDR based on the Q value cut-off alone does not yield corresponding PQ-FDR levels (i.e. a Q value of 0.05 does not specifically produce either 5% or 1% PQ-FDR). Modifying the MTC significance value cut-off so that it takes into account the experimental variability inherent within the replicates helps to produce a more tailored list of differentially expressed protein identifications whilst controlling for PQ-FDR. Also, compounding the same-same technique with another method of filtering, such as fold change cut-offs, can reduce the number of false positives included in the final dataset, further reducing the PQ-FDR [30,31].

In this research article, we have demonstrated a revised method for statistical analysis for shotgun proteomics datasets. The same-same method facilitates the construction of post analysis P value histograms and aids the researcher in choosing an appropriate statistical testing protocol for their analysis. We have shown that in the right circumstances, using BH Q value cut-offs derived from the same-same analysis yields a set of results that provide more significantly differentially expressed proteins from a given dataset, while also determining PQ-FDR at the experimental level. In the future, we hope to expand on this methodology so that it can be applied equally well to other quantitative proteomics data types, and also develop new tests to build onto the existing same-same architecture to further improve the statistical rigour for all shotgun proteomics results.

## 5. Acknowledgments

The authors acknowledge support from BioPlatforms Australia through the Australian Government’s National Collaborative Research Infrastructure Scheme, and the Australian Research Council through Discovery Project DP190103140. The authors declare no conflict of interest.

## 6. Author contributions

DCH and PAH contributed equally to the conceptualization and design of this study. DCH was responsible for writing the software described. DCH prepared the original draft of the manuscript, and PAH was responsible for revision and editing.

## Notes

### Competing Interest Statement

The authors have declared no competing interest.

### Summary of Updates

Updated with minor clarifications and corrections to the text only. No change to the figures, tables, data or results.

## References

1. Colquhoun, D. An investigation of the false discovery rate and the misinterpretation of p-values. R Soc Open Sci 2014, 1, 140216.

2. Diz, A.P., Carvajal-Rodriguez, A., Skibinski, D.O. Multiple hypothesis testing in proteomics: A strategy for experimental work. Mol Cell Proteomics 2011, 10, M110004374.

3. Karp, N.A., Lilley, K.S. Design and analysis issues in quantitative proteomics studies. Proteomics 2007, 7 42–50.

4. Sullivan, G.M., Feinn, R. Using effect size-or why the p value is not enough. J Grad Med Educ 2012, 4, 279–282.

5. Streiner, D.L., Norman, G.R. Correction for multiple testing: Is there a resolution? Chest 2011, 140, 16–18.

6. Bender, R., Lange, S. Adjusting for multiple testing--when and how? J Clin Epidemiol 2001, 54, 343–349.

7. Armstrong, R.A. When to use the bonferroni correction. Ophthalmic Physiol Opt 2014, 34, 502–508.

8. Choi, H., Nesvizhskii, A.I. False discovery rates and related statistical concepts in mass spectrometry-based proteomics. J Proteome Res 2008, 7, 47–50.

9. Benjamini, Y., Hochberg, Y. Controlling the false discovery rate: A practical and powerful approach to multiple testing. J Royal Stat Society, Series B 1995, 289–300.

10. Holm, S. A simple sequentially rejective multiple test procedure. Scandinavian journal of statistics 1979, 6, 65–70.

11. Benjamini, Y., Yekutieli, D. The control of the false discovery rate in multiple testing under dependency. Annals of statistics 2001, 29, 1165–1188.

12. Carvajal-Rodriguez, A., de Una-Alvarez, J. Assessing significance in high-throughput experiments by sequential goodness of fit and q-value estimation. PLoS One 2011, 6, e24700.

13. Pascovici, D., Handler, D.C., Wu, J.X., Haynes, P.A. Multiple testing corrections in quantitative proteomics: A useful but blunt tool. Proteomics 2016, 16, 2448–2453.

14. Noble, W.S. How does multiple testing correction work? Nat Biotechnol 2009, 27, 1135–1137.

15. The, M., Kall, L. Integrated identification and quantification error probabilities for shotgun proteomics. Mol Cell Proteomics 2019, 18, 561–570.

16. Gierlinski, M., Gastaldello, F., Barton, G.J. Proteus: An r package for downstream analysis of maxquant output. BioRxiv 2018, doi.org/doi10.1101/416511.

17. Wieczorek, S., Combes, F., Lazar, C., Gianetto, Q.G., Gatto, L., Dorffer, A., Hesse, A.M., Coute, Y., Ferro, M., Bruley, C., et al. Dapar & prostar: Software to perform statistical analyses in quantitative discovery proteomics. Bioinformatics 2017, 33, 135–136.

18. Goeminne, L.J.E., Gevaert, K., Clement, L. Experimental design and data-analysis in label-free quantitative lc/ms proteomics: A tutorial with msqrob. J Proteomics 2018, 171, 23–36.

19. Zhang, X., Smits, A.H., van Tilburg, G.B., Ovaa, H., Huber, W., Vermeulen, M. Proteome-wide identification of ubiquitin interactions using ubia-ms. Nat Protoc 2018, 13, 530–550.

20. Efstathiou, G., Antonakis, A.N., Pavlopoulos, G.A., Theodosiou, T., Divanach, P., Trudgian, D.C., Thomas, B., Papanikolaou, N., Aivaliotis, M., Acuto, O., et al. Proteosign: An end-user online differential proteomics statistical analysis platform. Nucleic Acids Res 2017, 45, W300–W306.

21. Singh, S., Hein, M.Y., Stewart, A.F. Msvolcano: A flexible web application for visualizing quantitative proteomics data. Proteomics 2016, 16, 2491–2494.

22. Strimmer, K. A unified approach to false discovery rate estimation. BMC Bioinformatics 2008, 9, 303.

23. Choi, M., Chang, C.Y., Clough, T., Broudy, D., Killeen, T., MacLean, B., Vitek, O. Msstats: An r package for statistical analysis of quantitative mass spectrometry-based proteomic experiments. Bioinformatics 2014, 30, 2524–2526.

24. Ritchie, M.E., Phipson, B., Wu, D., Hu, Y., Law, C.W., Shi, W., Smyth, G.K. Limma powers differential expression analyses for rna-sequencing and microarray studies. Nucleic Acids Res 2015, 43, e47.

25. Gelman, A., Hill, J., Yajima, M. Why we (usually) don’t have to worry about multiple comparisons. J Res Educ Effect 2012, 5, 189–211.

26. Kall, L., Storey, J.D., MacCoss, M.J., Noble, W.S. Posterior error probabilities and false discovery rates: Two sides of the same coin. J Proteome Res 2008, 7, 40–44.

27. Rothman, K.J. No adjustments are needed for multiple comparisons. Epidemiology 1990, 1, 43–46.

28. Gierlinski, M., Cole, C., Schofield, P., Schurch, N.J., Sherstnev, A., Singh, V., Wrobel, N., Gharbi, K., Simpson, G., Owen-Hughes, T., et al. Statistical models for rna-seq data derived from a two-condition 48-replicate experiment. Bioinformatics 2015, 31, 3625–3630.

29. Tusher, V.G., Tibshirani, R., Chu, G. Significance analysis of microarrays applied to the ionizing radiation response. Proc Natl Acad Sci U S A 2001, 98, 5116–5121.

30. Tyanova, S., Temu, T., Sinitcyn, P., Carlson, A., Hein, M.Y., Geiger, T., Mann, M., Cox, J. The perseus computational platform for comprehensive analysis of (prote)omics data. Nat Methods 2016, 13, 731–740.

31. Tyanova, S., Temu, T., Cox, J. The maxquant computational platform for mass spectrometry-based shotgun proteomics. Nat Protoc 2016, 11, 2301–2319.

32. Neilson, K.A., Ali, N.A., Muralidharan, S., Mirzaei, M., Mariani, M., Assadourian, G., Lee, A., van Sluyter, S.C., Haynes, P.A. Less label, more free: Approaches in label-free quantitative mass spectrometry. Proteomics 2011, 11, 535–553.

33. Neilson, K.A., Keighley, T., Pascovici, D., Cooke, B., Haynes, P.A. Label-free quantitative shotgun proteomics using normalized spectral abundance factors. Methods Mol Biol 2013, 1002, 205–222.

34. George, I.S., Pascovici, D., Mirzaei, M., Haynes, P.A. Quantitative proteomic analysis of cabernet sauvignon grape cells exposed to thermal stresses reveals alterations in sugar and phenylpropanoid metabolism. Proteomics 2015, 15, 3048– 3060.

35. Wu, Y., Mirzaei, M., Pascovici, D., Chick, J.M., Atwell, B.J., Haynes, P.A. Quantitative proteomic analysis of two different rice varieties reveals that drought tolerance is correlated with reduced abundance of photosynthetic machinery and increased abundance of clpd1 protease. J Proteomics 2016, 143, 73–82.

36. Mishra, P., Eich, M. Join processing in relational databases. CM Computing Surveys 1992, 24, 63–113.

37. Hommel, G., Krummenauer, F. Improvements and modifications of tarone’s multiple test procedure for discrete data. Biometrics 1998, 54, 673–681.

38. Qian, H.R., Huang, S. Comparison of false discovery rate methods in identifying genes with differential expression. Genomics 2005, 86, 495–503.

